# pholidosis: an R package to compare biological surface patterns

**DOI:** 10.1101/2025.02.23.639543

**Authors:** Isaac W Krone

**Affiliations:** Museum of Vertebrate Zoology, Dept of Integrative Biology, University of California, Berkeley

## Abstract

Atop the heads of many lizards and snakes sit complex patterns of flattened scales, differing in their sizes, shapes, colors, textures, and inter-relationships. These patterns of head scalation–called “pholidosis” – are often diagnostic to species or genera and therefore of great importance to reptile taxonomy, yet they have never been investigated in a detailed comparative context. Similar patterns can be seen in the scutes of turtle shells and the “cells” of insect wings; these each consist of a surface pattern made up of discrete units that can be considered homologous between individuals, populations, and species. Here, I describe an R package, `pholidosis`, designed to analyze these patterns as networks, providing tools for their construction, analysis, and visualization, and comparative analysis via an edit distance metric.

## Introduction

This paper briefly describes and demonstrates the use of the `pholidosis` package for working with and comparing biological surface patterns. The theoretical basis of this implementation is described in Krone (2025).

This paper has three main sections; the ‘Description’ section describes input data formats and functions of the `pholidosis` package, briefly explaining their structure and use and outlining cases for their use. The ‘Vignette’ section provides a worked example demonstrating the use of the R package with a dataset distributed with the package (modified from Krone (2025)). Finally, the ‘Troubleshooting’ section describes methods to visualize and analyze input data that cause errors or warnings, and how to go about solving the issues.

## Description

### Input data

The basic currency of the `pholidosis` package is the pholidosis network (Krone, 2025), implemented as an igraph object (Csárdi et al., 2025; Csárdi & Nepusz, 2006).

Constructing a network can be accomplished in many ways, but the `pholidosis` package is built to easily construct networks from adjacency matrices coded into an .xlsx spreadsheet. There are several advantages to working with a spreadsheet of adjacency matrices as a basic data input structure. First, all of the matrices, and therefore all of the network data needed to perform an analysis, can be stored in a single file readable by most commonly-used office suites. This allows for easy data sharing, as well as ease of copying of and comparisons between the individual adjacency matrices. Second, an adjacency matrix clearly lists all of the individual nodes in a network and presents their adjacency relationships in a human-readable format. Finally, conditional formatting and macros within spreadsheet software can substantially improve readability of adjacency matrices and reduce the time needed to construct them by copying large chunks of the matrix (for instance, the connections of the right and left sides of a symmetrical structure).

Adjacency matrices for the current version of `pholidosis` must have the following properties:

1. The first cell of the spreadsheet (A1) must be labelled “scale name”
2. The subsequent cells of row 1 and column A must contain unique names of all vertices in the network. Names of vertices not in the network may be included if, for instance, the spreadsheet is copied from a template that includes more vertices.
3. If vertices belong to a named group of vertexes (for instance, the supralabial scales of a squamate), they should be named according to the group, followed by an underscore and a number: for instance, supralabial_1.
4. Names of vertices should only begin with “R_” and “L_” if they belong to the right and left sides of a network, respectively.
5. Adjacency relationships between vertices are indicated by the numbers 1-3. The number 1 indicates two elements that are adjacent but fully separated or distinguished from each other via a clear boundary. The number 3 refers to two elements that are adjacent and fused together or have no clear boundary. The number 2 indicates two elements that are adjacent and somewhat fused or partially separated by an incomplete or indistinct boundary.
6. All cells other than those indicating adjacency should be left blank.

`pholidosis` provides an easy-to-use function, `excel_to_network()`, to read a spreadsheet workbook into R as a list of networks using the `readxl` library (Wickham et al., 2023). When run with the argument `“verbose = TRUE”`, `excel_to_network()`calls the function `verifyMatrix()` to provide context on problems with specific matrices, including unmatched row and column names, and non-numeric entries in the matrix.

For importing a network from a data frame, use the `scaleNetwork()` function, for which `excel_to_network()` is a wrapper. Note that `scaleNetwork()` changes edge weights; edges weighted 2 are changed to a weight of 1.5 and edges weighted 3 are changed to a weight of 2. This preserves the equivalency between fusion and loss of scales (Krone, 2025).

### Visualizing pholidosis networks

Tools to quickly plot network sand their properties are essential for communicating results and troubleshooting data. However, plotting networks in R is very much an art. Though pholidosis networks are by definition planar (meaning they should be able to be plotted two dimensionaly without overlapping edges), automated network plotting tools usually induce some distortion or overlap in pholidosis networks including more than a couple dozen vertices.

Therefore, `pholidosis` provides two tools to aid in plotting networks. The first, `lizard_setup()`, provides a setup function that helps to lay out the network in two dimensional space. It provides basic ordination for each vertex in the network by designating one vertex as at the extreme end of the network using the firstscale argument and measuring the distance of each vertex from it. It also “sides” scales by designating scale names beginning with “L_” as side -1 and scales beginning with “R_” as side 1. `lizard_setup()` is called by default by `scaleNetwork()`, but users can write their own setup functions to provide or modify network attributes on import via `scaleNetwork()` and `excel_to_network()`.

The second plotting tool, `pholidosis_plot()`, provides a quick method to plot a pholidosis networks set up by `lizard_setup()` using ggraph (Pedersen, 2024). By default, `pholidosis_plot()` plots the network using the igraph “stress” layout. It plots each type of vertex (identified via the `vertextype` attribute) as a different color and labels vertices only by any numbers in their name, allowing the positions of different vertices of the same vertextype to be distinguished by their number.

### Comparing pholidosis networks

The main point of the `pholidosis` package is to compare pholidosis networks. It accomplishes this by implementing a graph edit distance measurement. `pholidosis` determines the minimum number of changes–additions or subtractions of vertices (*f* distance), rearrangements of edges (*e* distance), and changes in edge weight (*w* distance)– necessary to transform one graph into the other and vice versa. This is accomplished via the `pnet_distance()` function, which calls `graph_edit_distance()` to determine the *f* and *e* distances and then calculates the *w* distance. The algorithm used by `graph_edit_distance()` is described in Krone(2025).

`pnet_distance()`compares pairs of networks, and can be slow depending on network complexity and dissimilarity. When troubleshooting networks that may have incorrect homologies or edges, it is useful to use `pnet_distance()` with the truncate=FALSE argument to retain full edge-by-edge comparison data frames generated by `graph_edit_distance()`.

To produce a distance matrix for a list of networks, use the `net_dist_mat()` function, which measures all three distances between every pair of networks in the list to produce an *n* × *n* × 3 matrix, where n is the number of networks in the list. Since computation takes some time, `net_dist_mat()` prints the pair of graphs `pnet_distance()`is comparing to the console as it runs. The network distance matrix allows for comparisons of network distance and any other trait between any pair of networks, forming the basis of comparative biology with this method. The *w, e* and *f* distances measure equivalent units (see Krone -Krone (2025)), and can therefore be summed to produce a composite network distance. An *n* × *n* × 1 composite network distance matrix provides a single metric for comparison between networks, which can be visualized using principal coordinates analysis and used to build distance-based phylogenetic hypotheses.

Another method of comparison between networks is character-based, rather than distance-based. The `morpho_matrix()` function produces a character matrix for a list of networks, using all edges found in all networks as potential characters and scoring edges as either absent (0) or as their edge weight. This character matrix can be used to build character-based phylogenetic hypotheses.

### Vignettes

This series of vingettes provides several examples of how to use the `pholidosis` package. The first vignette covers installation, importing adjacency matrices, visualizing pholidosis networks, creating an edit distance matrix, and producing phylogenetic hypotheses using pholidosis network data. The second vignette covers troubleshooting procedures for use when importing and comparing networks.

### Workflow

#### Installation

`pholidosis` makes heavy use of the `igraph` R package (Csárdi et al., 2025; Csárdi & Nepusz, 2006). See https://r.igraph.org/ for instructions on installing this package, which may require installing dependencies outside of R.

`pholidosis` can be installed from GitHub with:

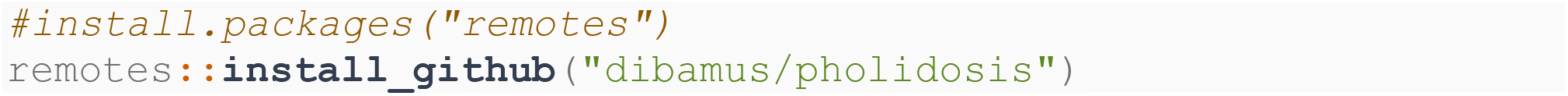

#### Import

Import the `pholidosis` library.

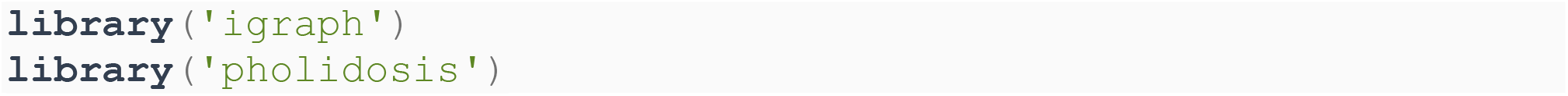

`pholidosis` includes a demo dataset of six pholidosis networks of dibamid lizards from Krone -Krone (2025). This is stored as an excel file (.xlsx), so you can inspect the file in a spreadsheet editor to better understand how to set up an adjacency matrix for `pholidosis`.

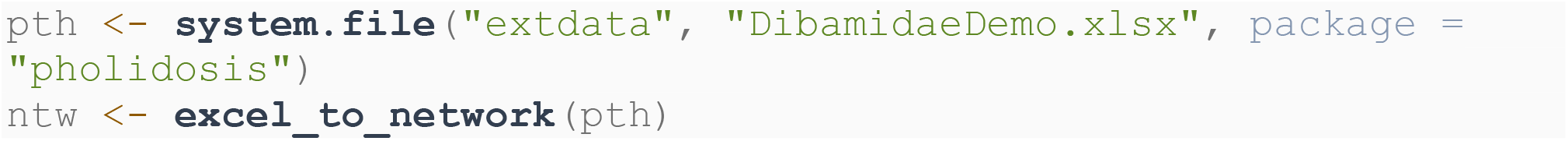

#### Visualization

After loading the dataset, we can quickly visualize the networks via the `pholidosis_plot` function, which uses the `ggraph` library (Pedersen, 2024) to produce a `ggplot2` graph object (Wickham, 2016).

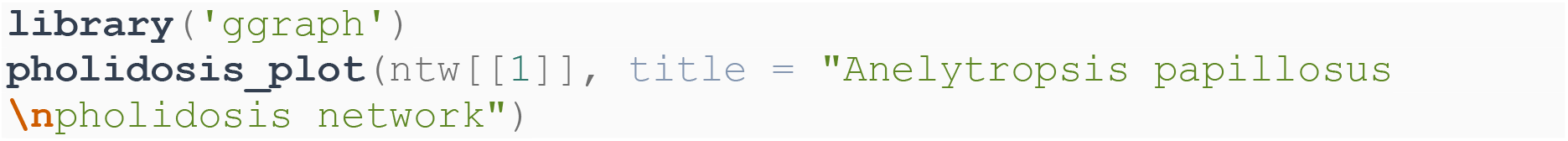

**Figure 1.**
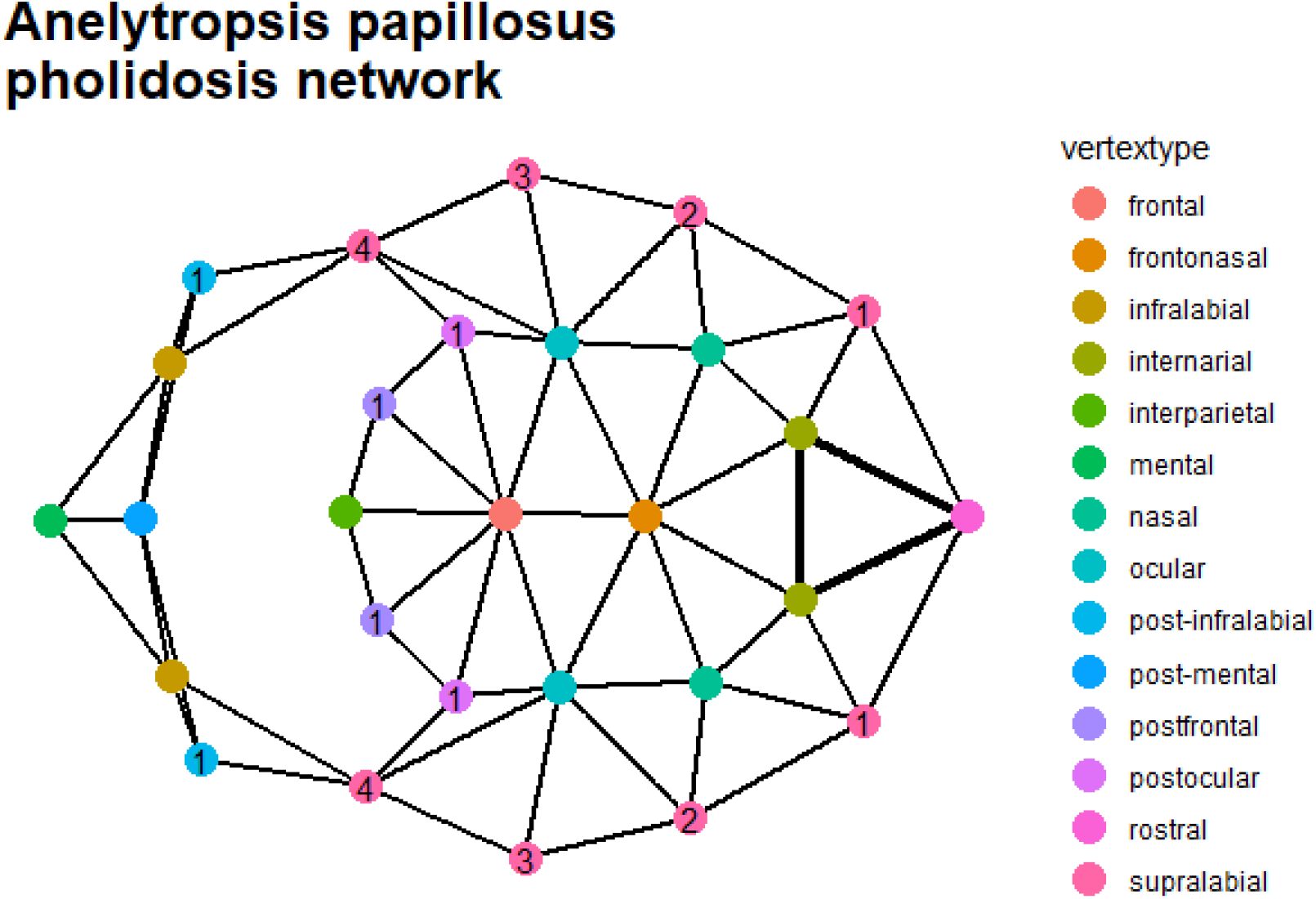
a pholidosis network plotted with pholidosis_plot()

In this visualization, we can see the topology of the pholidosis network laid flat, as if you were viewing the lizard’s head from above. The front of the lizard’s head is on the right side, and the rest of the lead is laid out towards the left. Vertices corresponding to the scales of the lower jaw are laid out further to the left. Here, vertices are color coded by the scale type they represent. Numbers appear in vertices where those vertices represent the same type of scale (for instance, a supralabial). The bold lines between the rostral and internarial vertices indicate a higher edge weight; those scales are fused together in this animal.

Note that the automatic layout method (`‘stress’`) isn’t perfect; graphs will often be “stretched” or “folded” rather than laid out totally flat. You can see this in the plotting of the infralabial and post-infralabial vertices.

We can also check on a few basic properties of pholidosis networks:

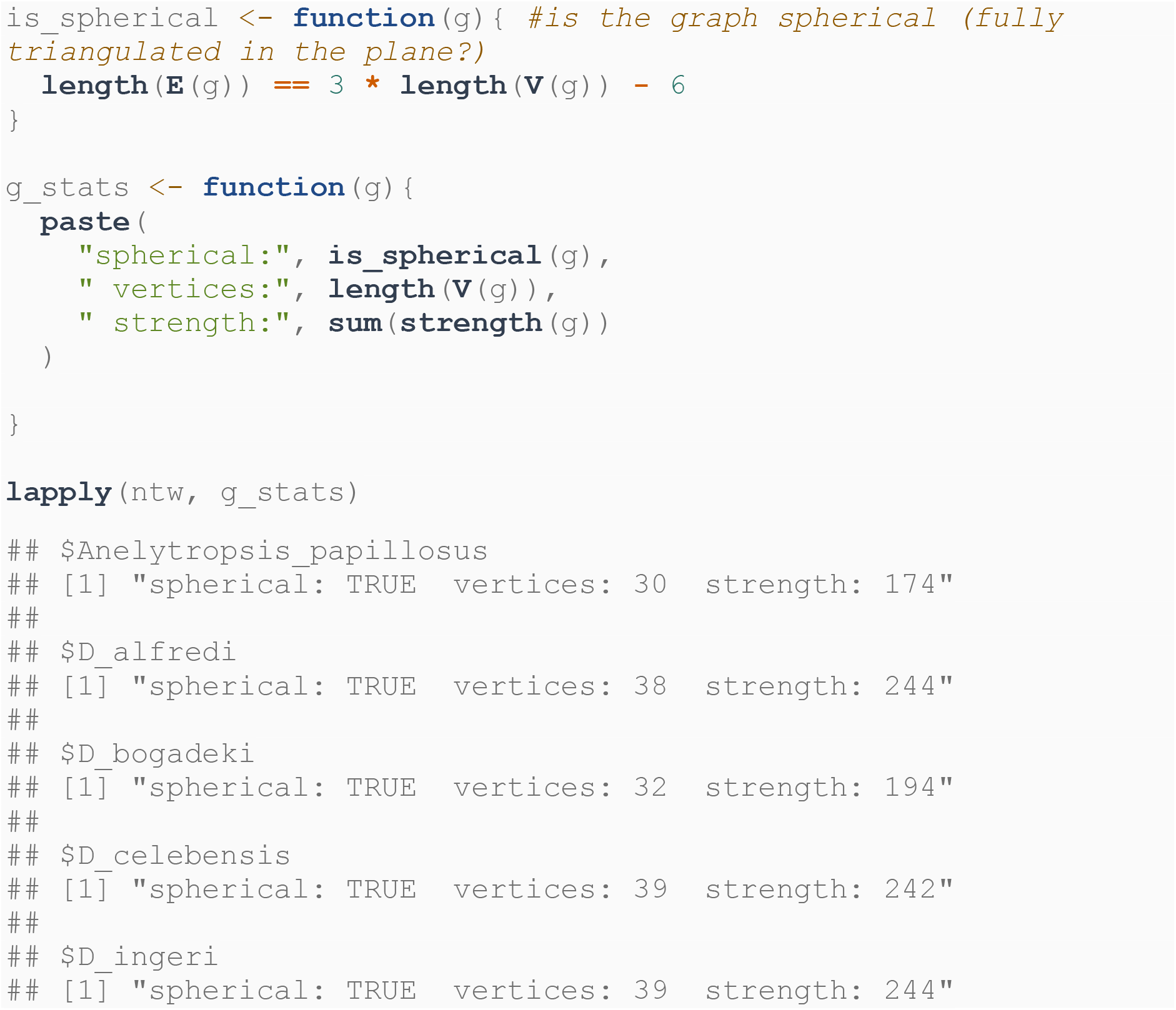
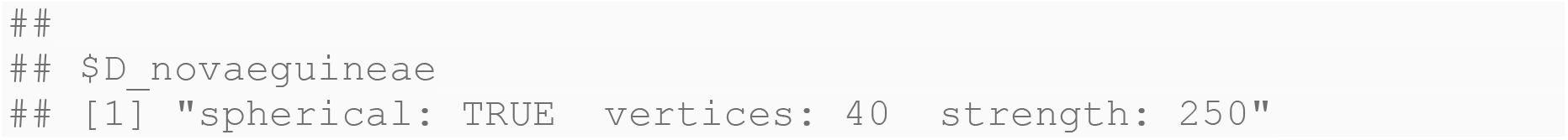

#### Comparison

`pholidosis` implements an edit-distance metric for pholidosis networks, allowing for consistent comparisons between the networks. Any two networks can be compared via the function `pnet_distance()`. Calling `pnet_distance()` on a pair of graphs returns the *w, e*,and *f* distances.

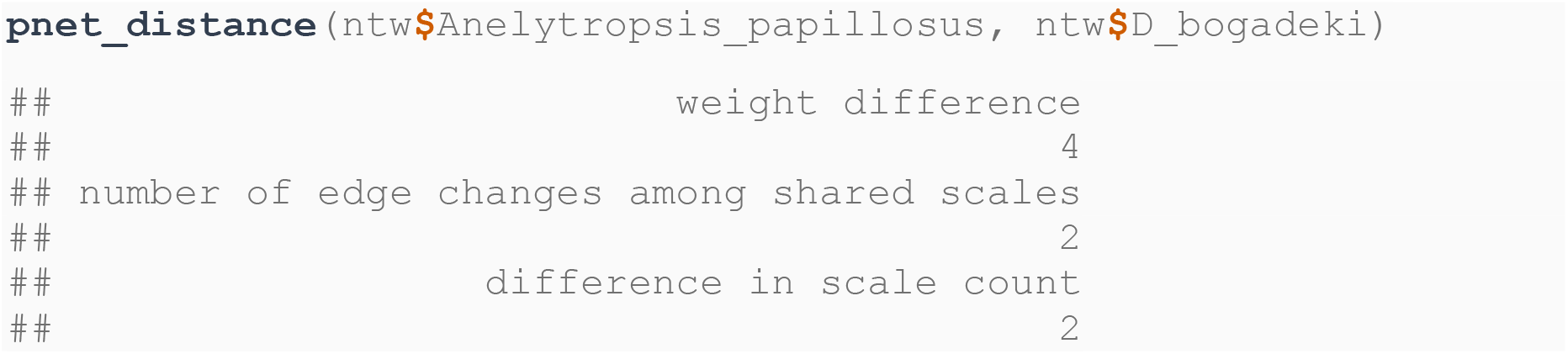

Adding the argument `truncate = FALSE` to the call returns data frames enumerating the exact changes needed to change each graph into the other.

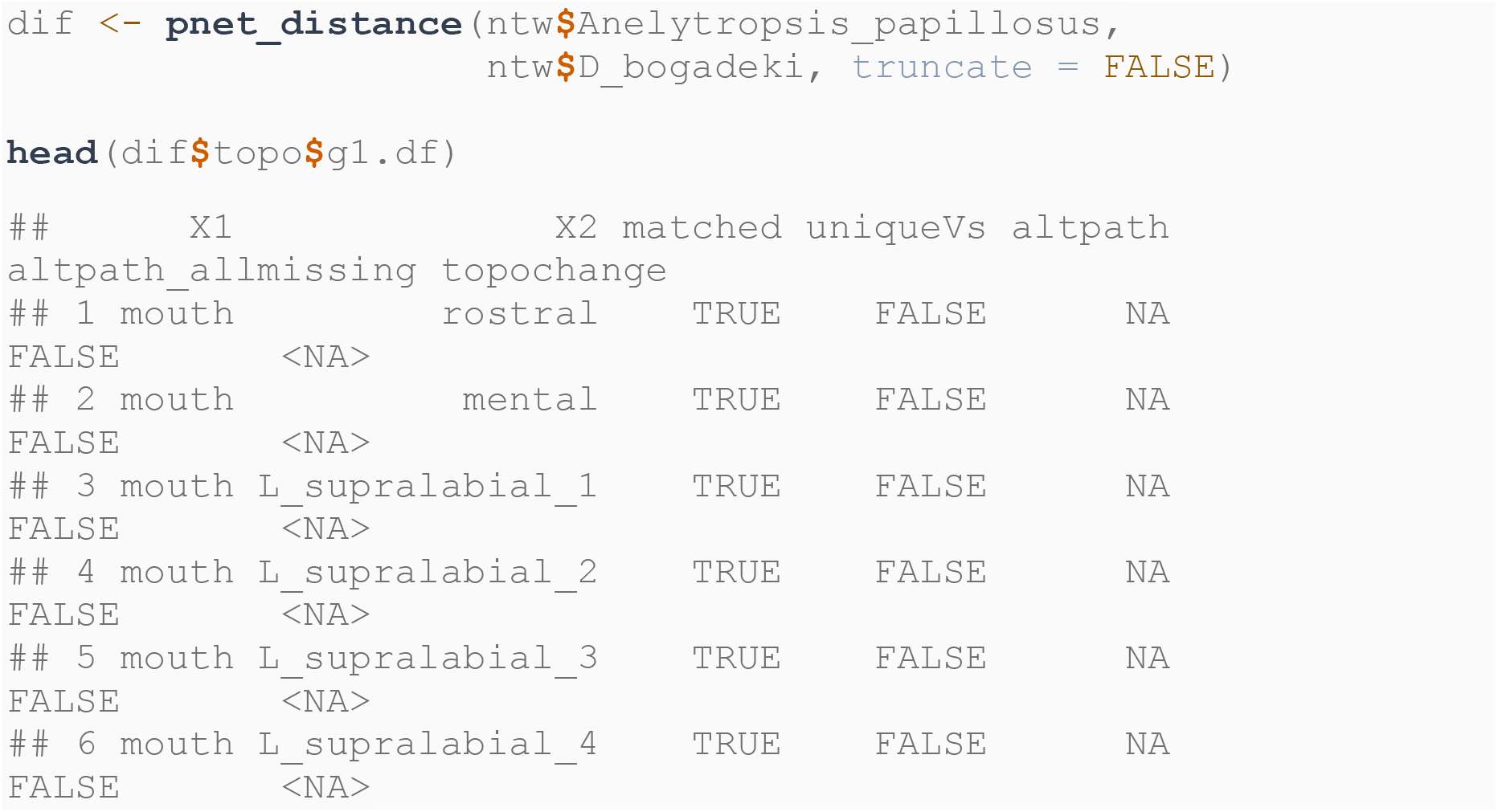
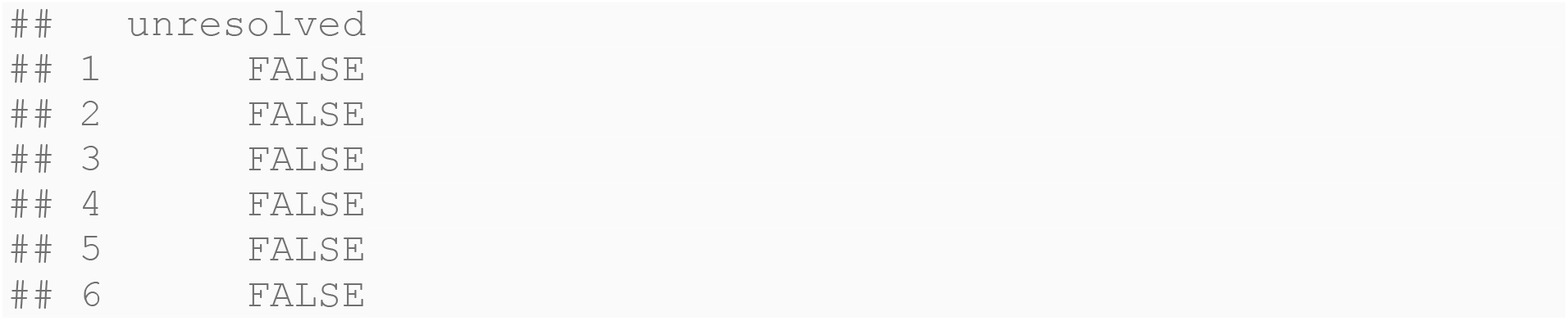

The `net_dist_mat` function provides a quick way to produce matrices of edit distances. Producing an edit-distance matrix is computationally intensive: the following chunk may take a minute or so to run.

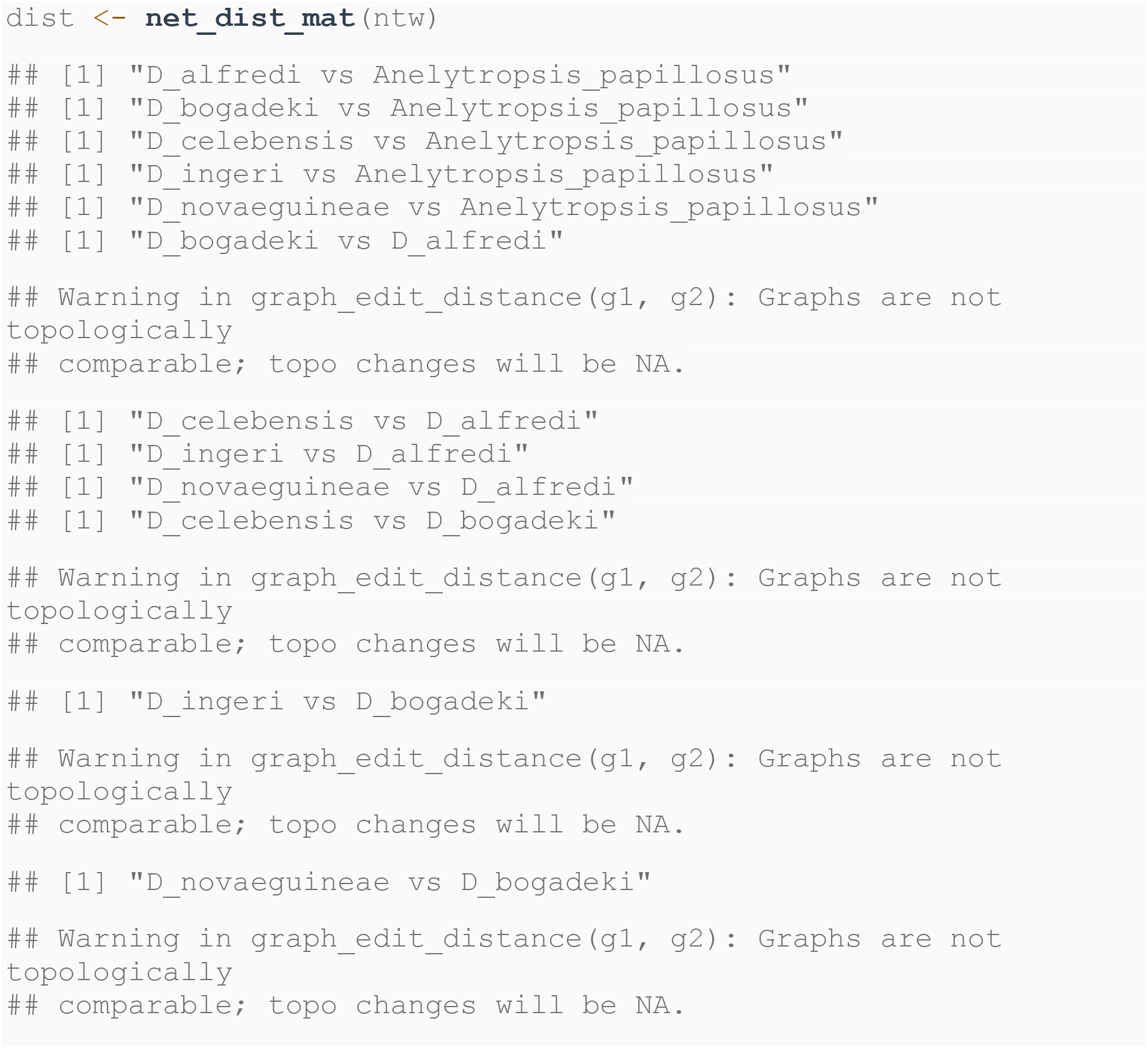
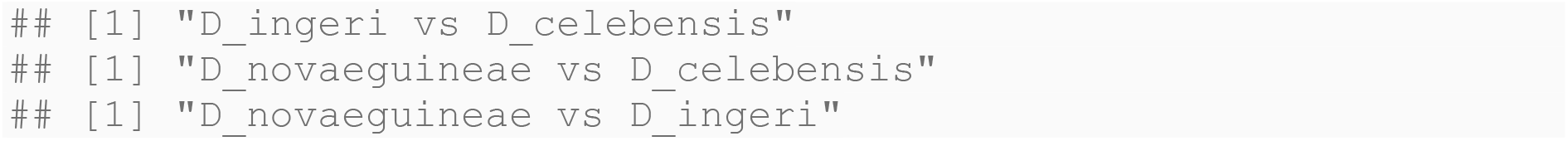

Note that the matrix is three dimensional, with a size of *n* × *n* × 3, where n is the length of the list of graphs input. There is an *n* × *n* matrix for each of the 3 measured distance components.

Separating out these components can help you understand how these different features are contributing to the diversity of the system you’re studying, but the measurements contained in them are equivalent. For instance, we can see that only *Dibamus booliati* has edge rearrangements relative to the other taxa.

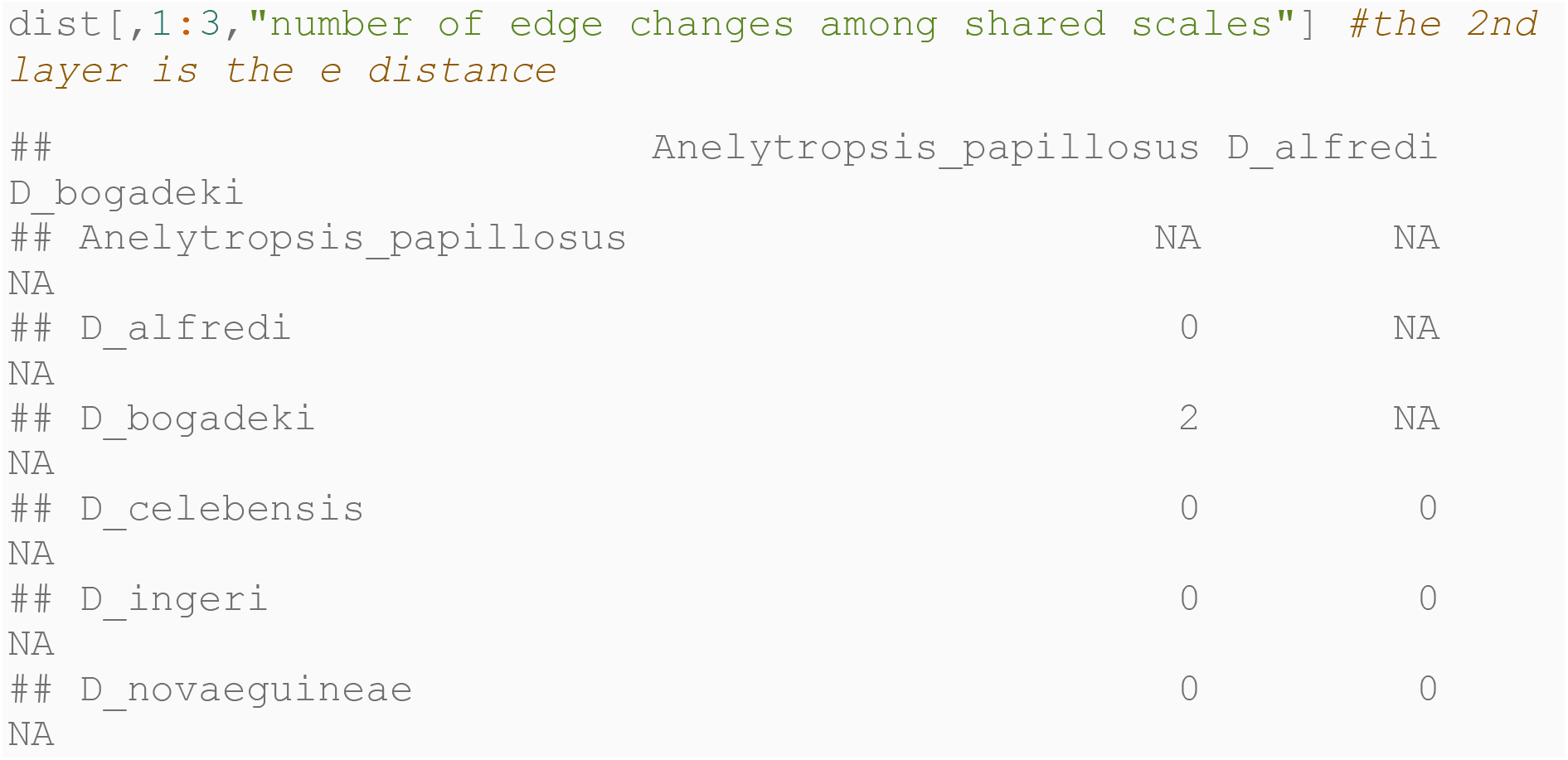

If you want to continue with analysis, combine these 3 components into a single matrix.

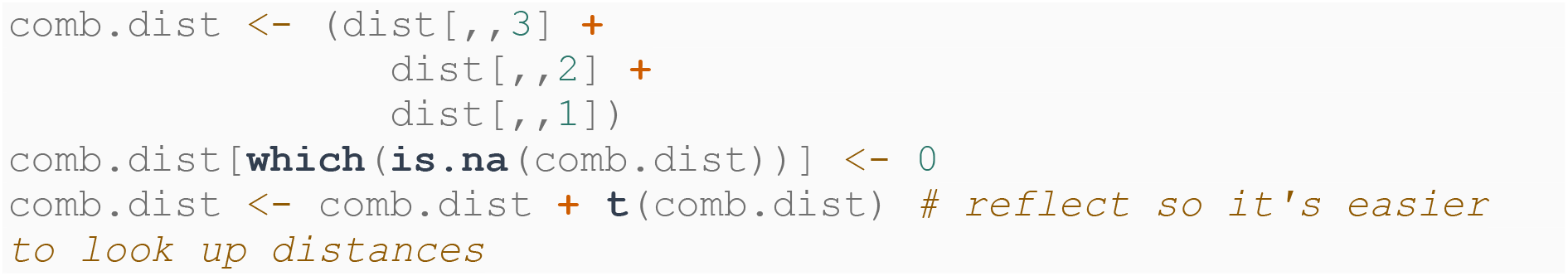

You can easily visualize the distances by visualizing the matrix as a heatmap with ggplot (Wickham, 2016).

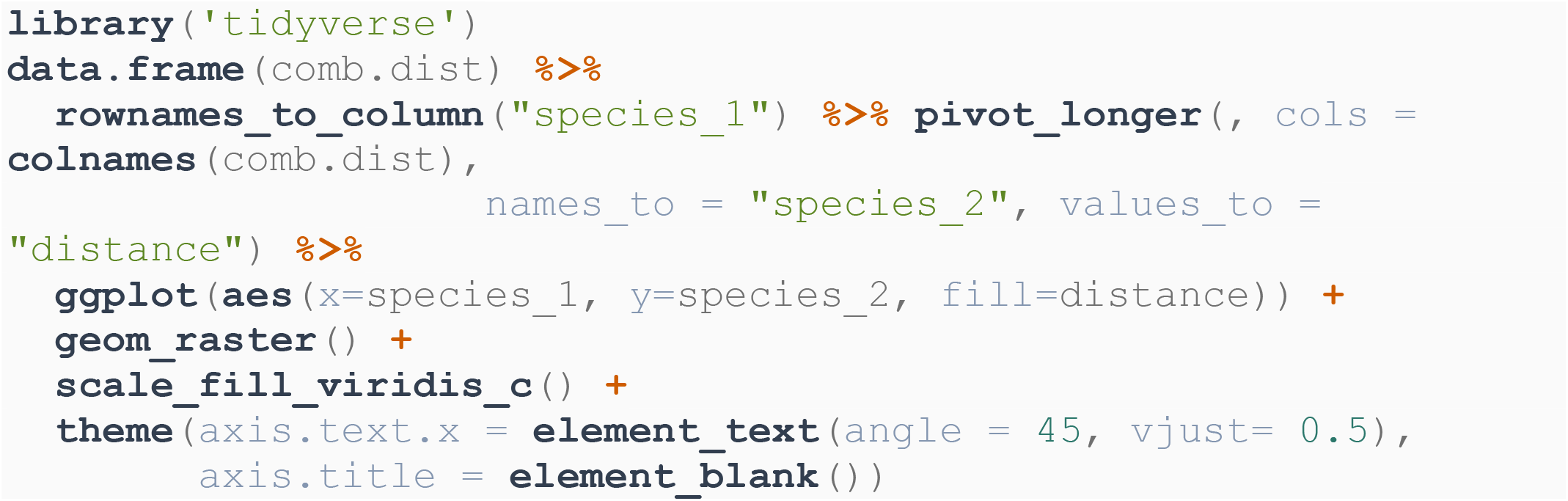

**Figure 2.**
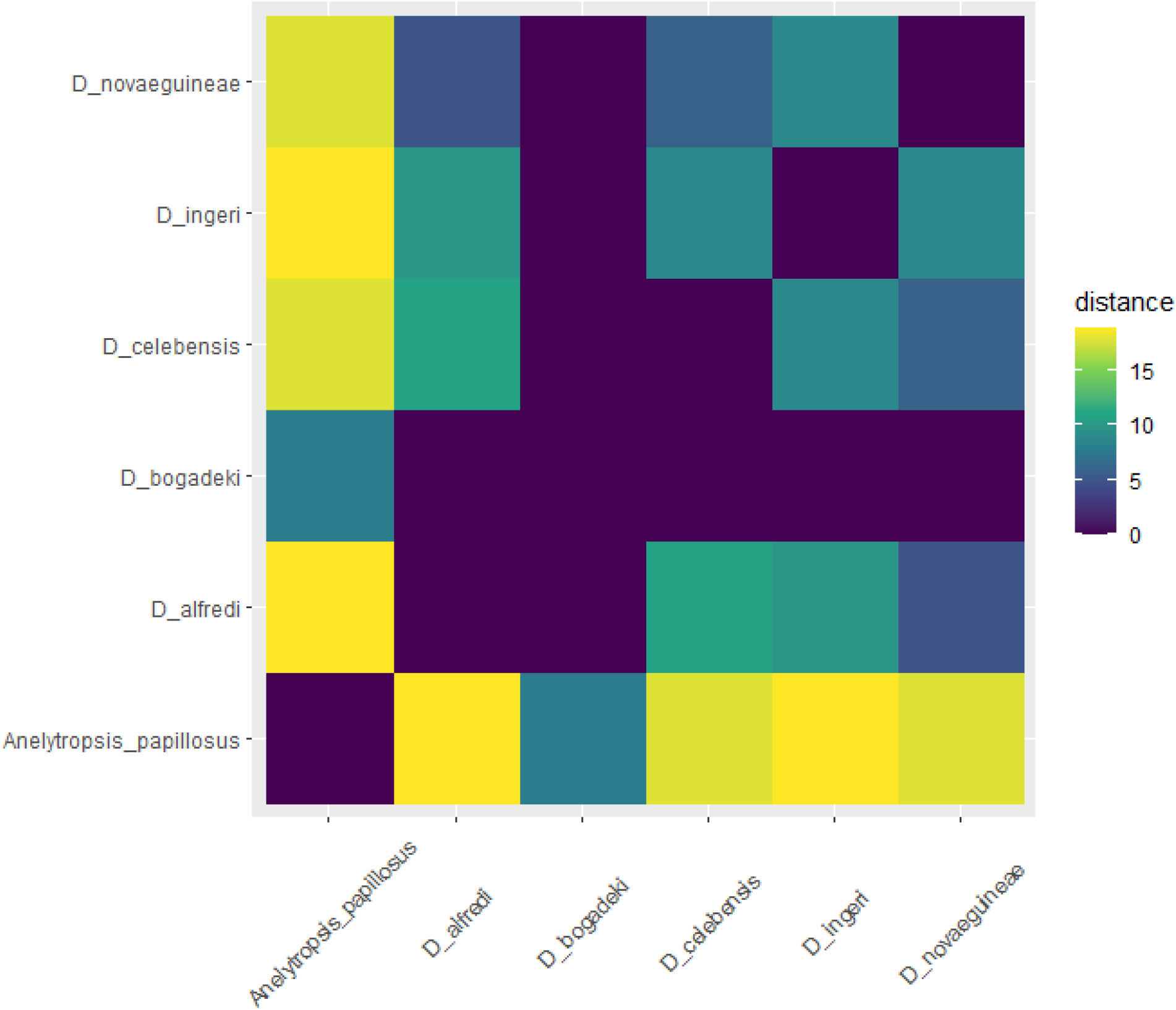
Combined distances of dibamids demo networks.

Principal Coordiantes analysis can reduce this to two dimensions

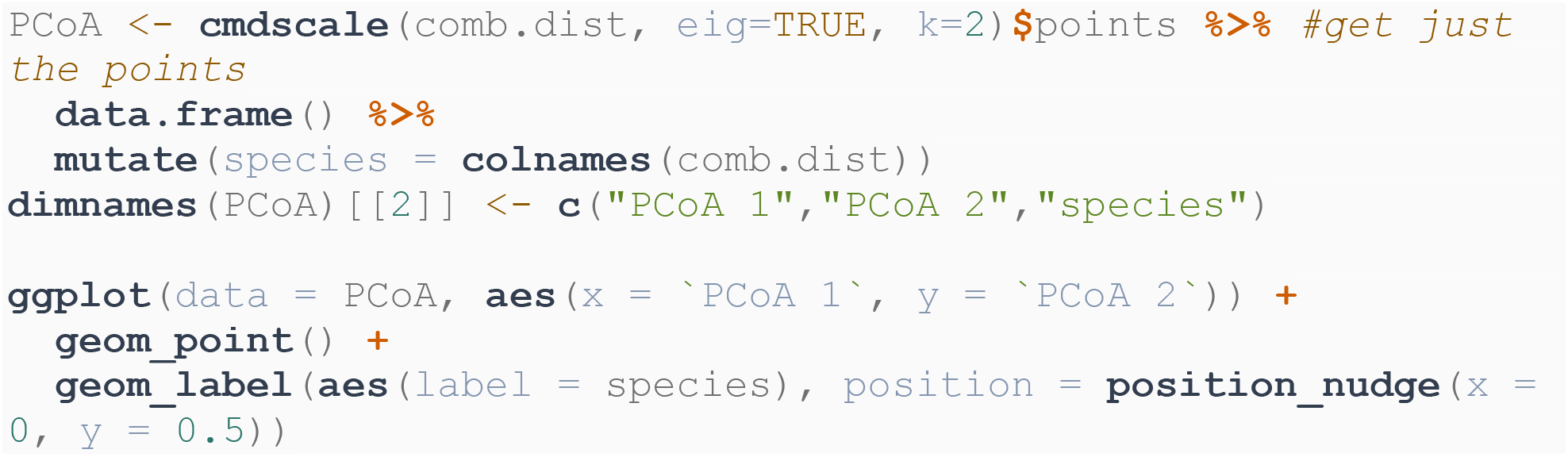

**Figure 3.**
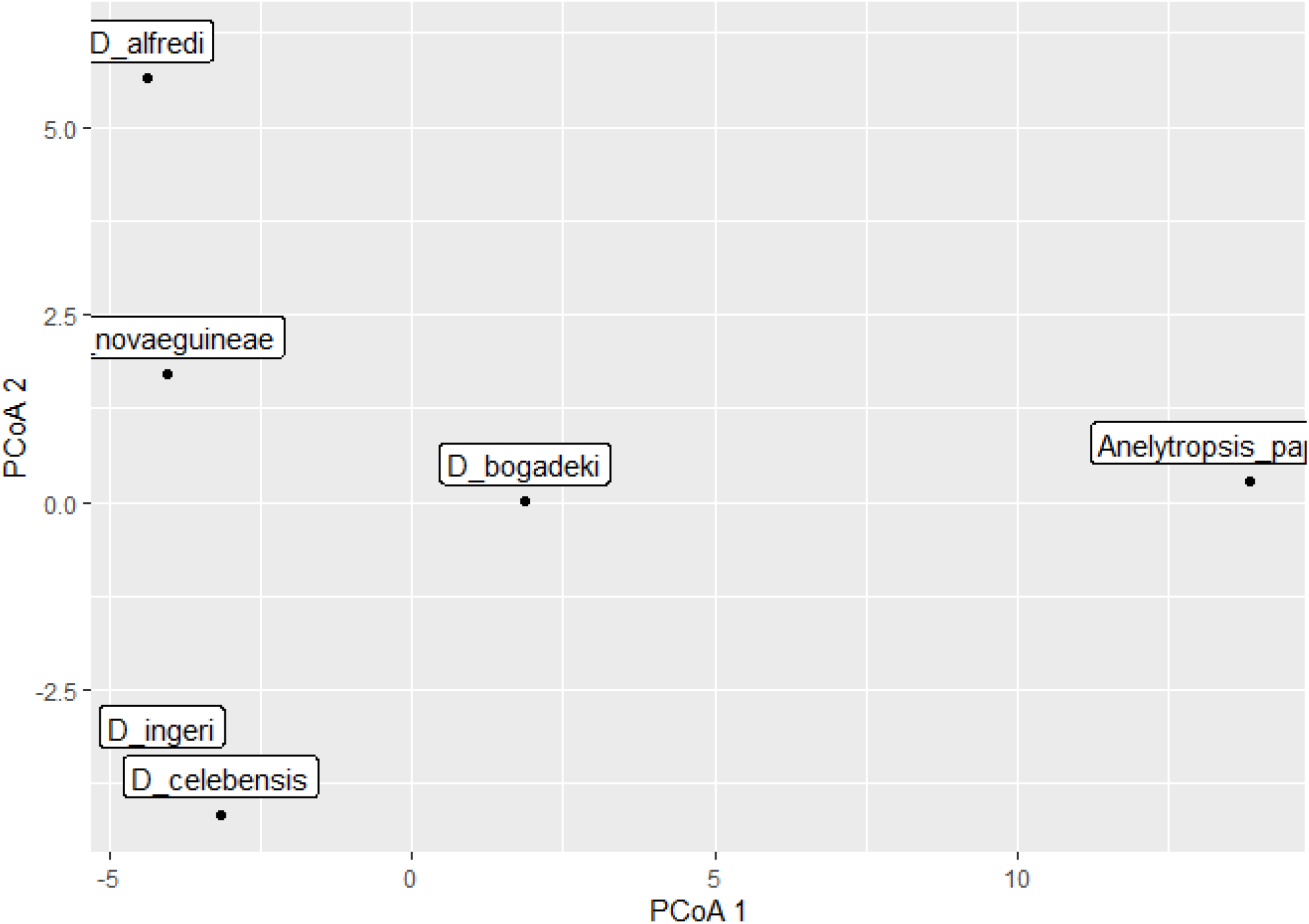
Principal coordinates analysis of dibamids demo networks.

#### Phylogenetics

Pholidosis networks can be used to produce or inform phylogenetic hypotheses. Distance-based phylogenies can be quickly constructed based on the distance matrix via the `nj` function in `ape` (Paradis & Schliep, 2019).

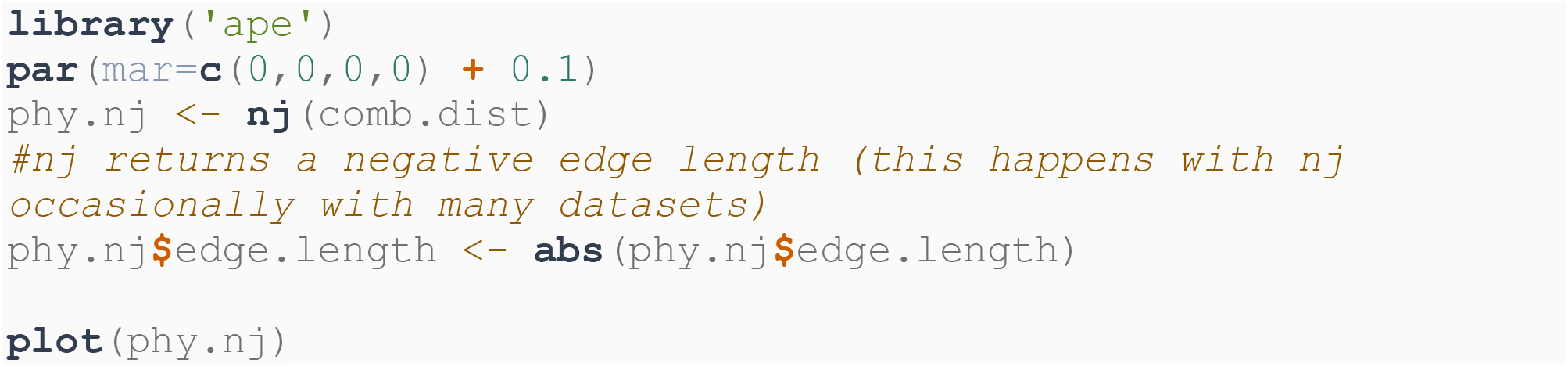

**Figure 4.**
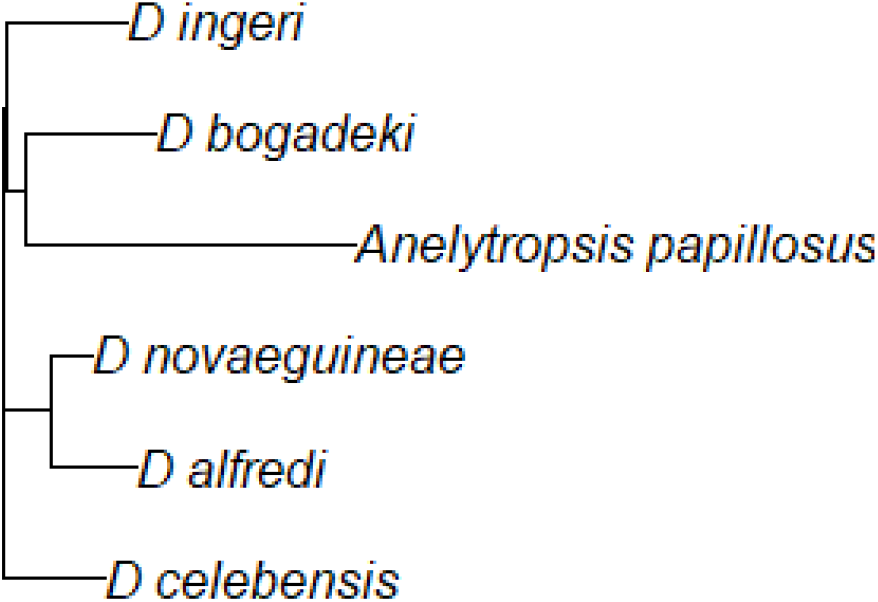
Distance-based phylogeny of dibamids demo networks.

Alternatively, `pholidosis` implements a function to produce a character matrix based on a list of pholidosis networks. The characters involved are edges, their states are the edge weights. We can use the `pratechet` function in `phangorn` (Schliep, 2011) to reconstruct a phylogenetic tree using the parsimony ratchet.

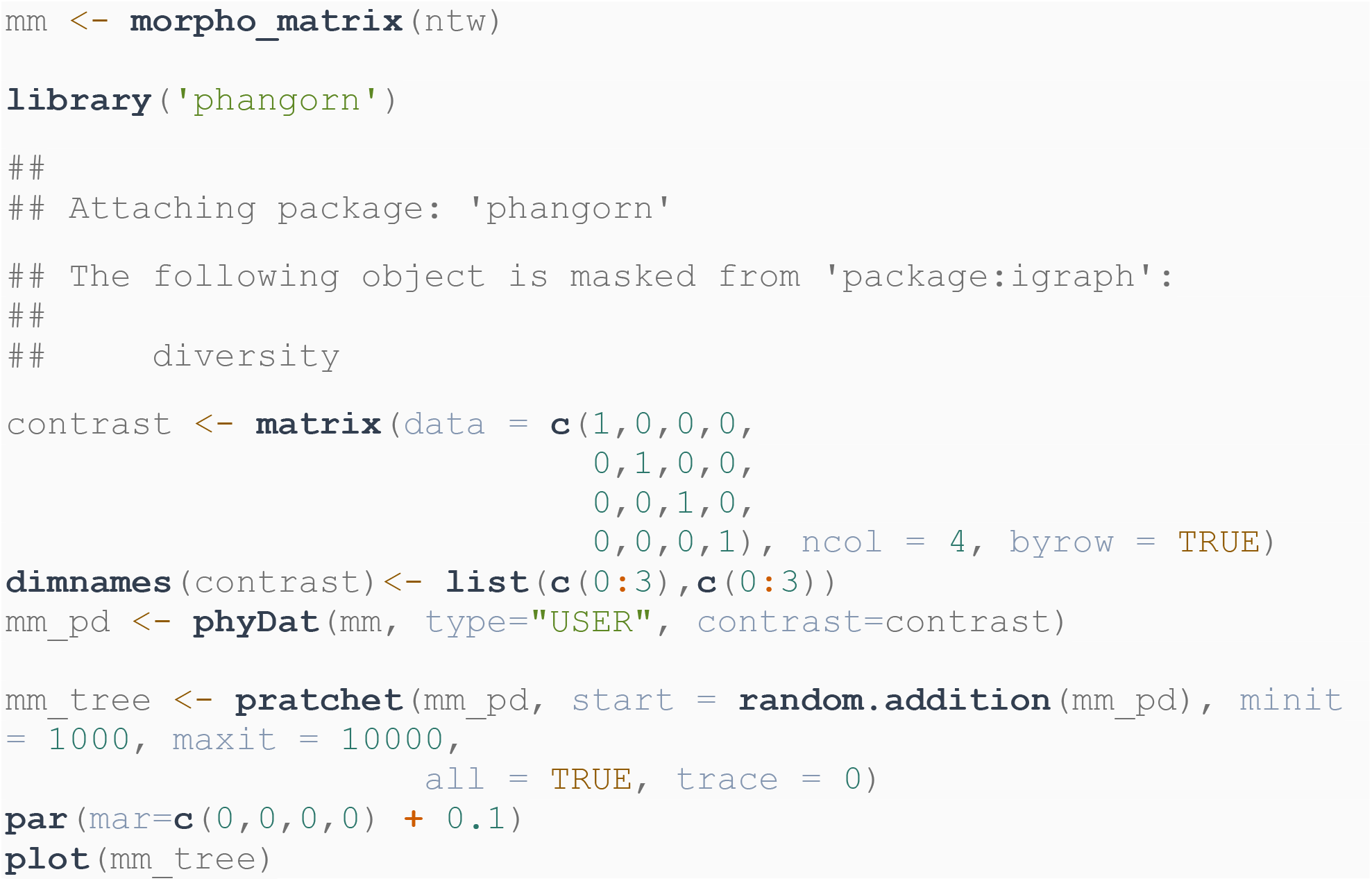

**Figure 5.**
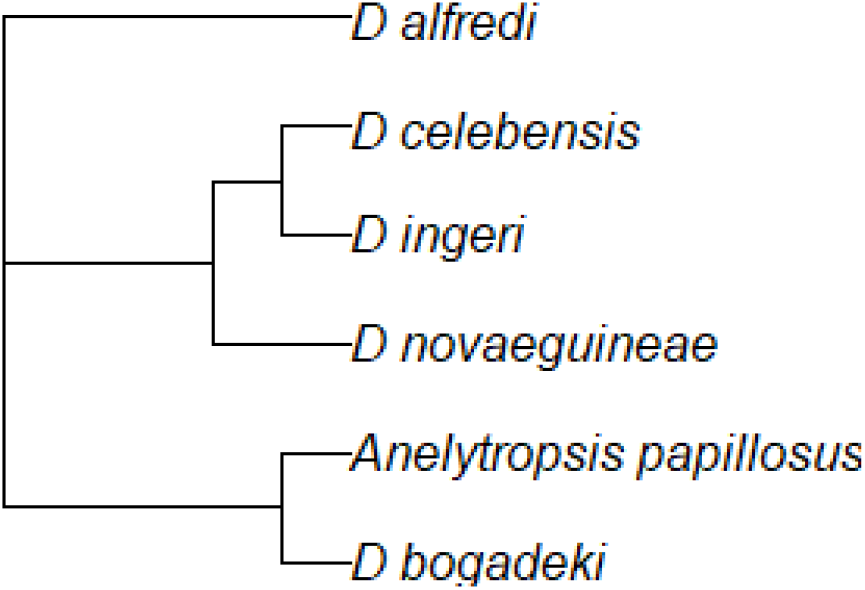
Distance-based phylogeny of dibamids demo networks.

#### Troubleshooting

The most difficult part of an analysis using pholidosis networks is collecting and correctly formatting the input data. The `pholidosis` functions for importing matrix data are designed to catch many small errors in input matrices to aid in troubleshooting new datasets. It also includes a few example datasets containing common mistakes.

Try to import the matrices in “bad_matrix.xlsx”

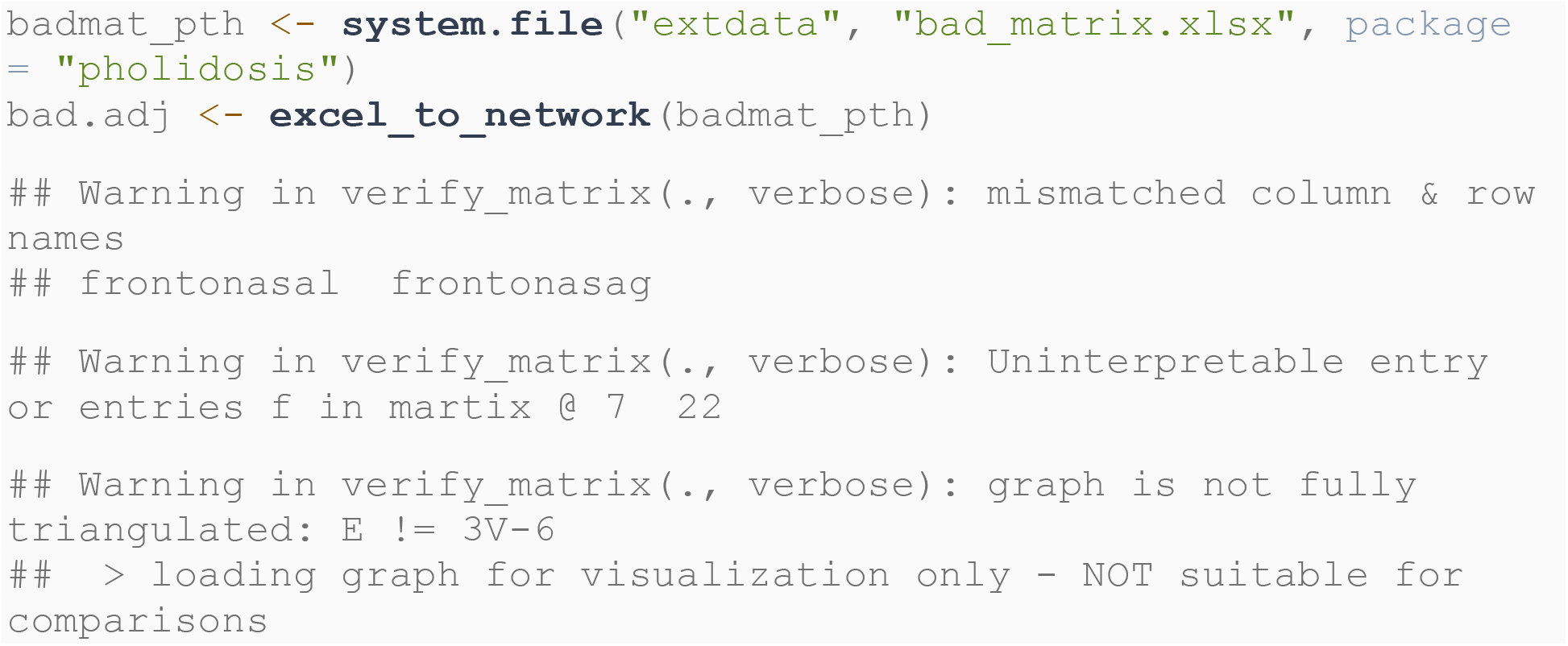

Quite a few warning messages there. To get more information on why it didn’t work, we can call `excel_to_network` with the argument `verbose = TRUE`.

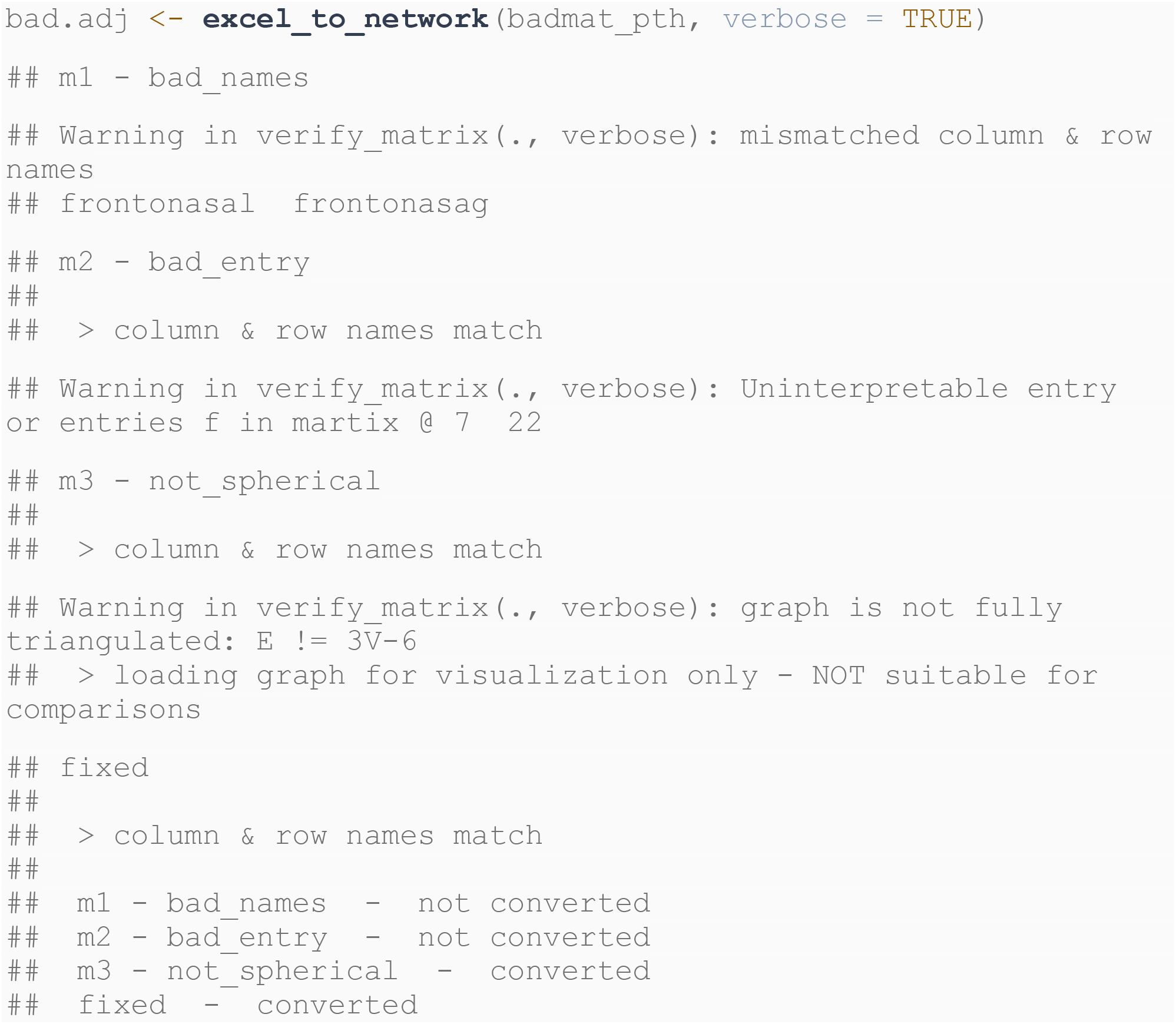

When `verbose = TRUE`, the function narrates what sheets it’s reading and when it determines whether column & row names are matching, then produces a table noting whether sheets were converted into networks.

What are the errors here?

- The first sheet was not converted because of mismatched row and column names; the column “frontonasag” is misspelled: it should be “frontonasal.”
- The second sheet was not converted because there was an uninterpretable input in the sheet - the cell at `[7,22]` contains `f`, rather than an interpretable value (NA and numbers 0,1,2 & 3)
- The third sheet was converted, but it is topologically invalid. All pholidosis networks must have 3*V* − 6 edges (where *V* is the number of vertices) to be fully triangulated. This sheet was converted into a network, but it should not be compared to any other graph via `graph_edit_distance`, since it does not comply with the topological constraint of full triangulation. Plotting it might show missing or excess edges.

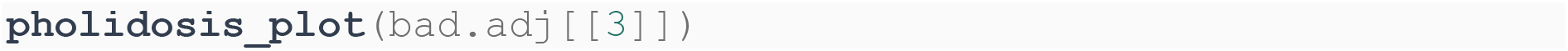

**Figure 6.**
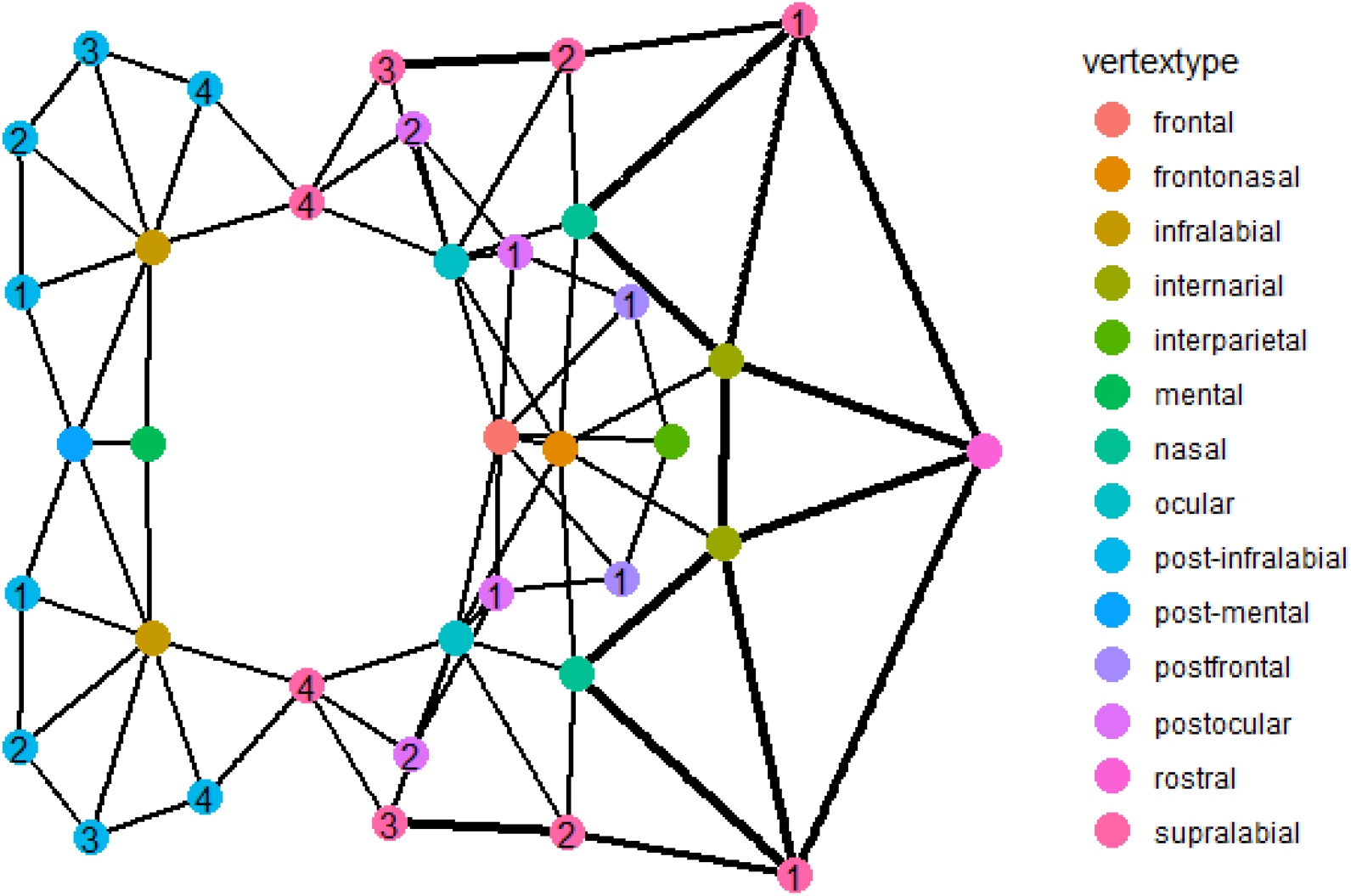
a problematic pholidosis network plotted with pholidosis_plot()

Nothing appears to be wrong with it. But this is illusory; it is missing “mouth” and “body” vertices, which are by default not plotted by `pholidosis_plot()`.

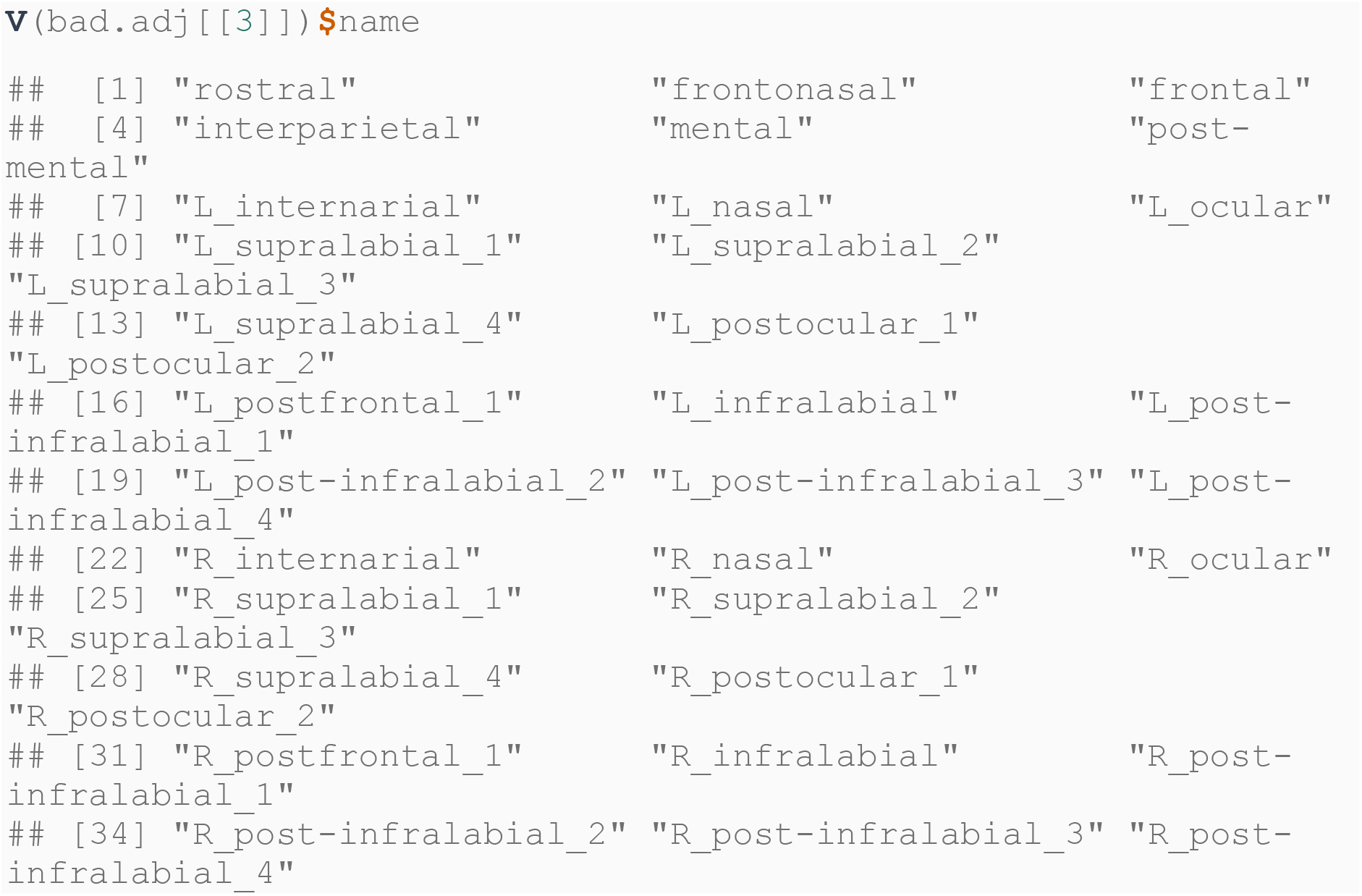

Most problems encountered when using `pholidosis` stem from bad input graphs, so users are advised to plot out graphs after you import to try to catch early mistakes.

## Acknowledgements

Thanks to Dr. Ian Wang for encouraging me to produce this publication.

